# Pathogen transmission modes determine contact network structure, altering other pathogen characteristics

**DOI:** 10.1101/2022.08.25.505277

**Authors:** Melissa Collier, Gregory F Albery, Grant C. McDonald, Shweta Bansal

## Abstract

Pathogen traits can vary greatly and impact the ability of a pathogen to persist in a population. Although this variation is fundamental to disease ecology, little is known about the evolutionary pressures that drive these differences, particularly where they interact with host behavior. We hypothesized that host behaviors relevant to different transmission routes give rise to differences in contact network structure, constraining the space over which pathogen traits can evolve to maximize fitness. Our analysis of 232 contact networks across mammals, birds, reptiles, amphibians, arthropods, fish, and mollusks found that contact network topology varies by contact events, most notably in networks that are representative of fluid-exchange transmission. Using infectious disease model simulations, we showed that these differences in network structure suggest pathogens transmitted through fluid-exchange contact events will need traits associated with high transmissibility to successfully proliferate, compared to pathogens that transmit through other types of contact. These findings were supported through a review of known traits of pathogens that transmit in humans. Our work demonstrates that contact network structure may drive the evolution of compensatory pathogen traits according to transmission strategy, providing essential context for understanding pathogen evolution and ecology.

## 1 Introduction

Pathogens vary in a range of important characteristics including transmission mode, infectivity, and duration of infection, many of which determine epidemiological characteristics such as their ability to persist in a population (1–5). Although this diversity in pathogen traits is fundamental to disease ecology, we know little about the ecological factors driving the evolution of such traits; in particular, it is unclear how transmission ecology determines the evolution of pathogen characteristics.

Pathogens are spread by a range of different contact events facilitated by specific host behaviors such as respiration, physical contact, or shared space use, which define different pathogen transmission modes (6). In a contact network, the behavior that defines its edges (i.e. a contact event) can be associated with different transmission modes (e.g. mating vs. grooming vs. spatial associations), and exhibit distinctive contact patterns (7–11). For instance, when analyzing contact events in mice (*Mus musculus*), researchers found that agonistic, grooming, and sniffing events were associated with distinct network properties such as density, average path length, and node centrality (12). Such network properties can influence the transmission efficiency of pathogens, with downstream impacts on the evolution of their traits (13–19). Furthermore, it is known that individual contact effort across contact events can be heterogeneous (a concept known as social fluidity), and can lead to the formation of weak ties (9). These weak ties play an important role in defining network structure (18), but the extent to which they impact the evolution of pathogen traits remains unknown. Understanding variation in these characteristics is essential for understanding pathogen ecology, and therefore for developing control measures and testing hypotheses regarding their evolutionary origins.

Pathogens should evolve to maximize their fitness, principally described by their *R*_0_ (i.e. the total number of new infections caused by one infection in a totally susceptible population) (20). To persist in a population, a pathogen’s *R*_0_ must be greater than 1. A pathogen’s *R*_0_ depends on both the behavior of its host population, and on its own transmissibility (1,21,22). Host behaviors create the relevant contact that defines the path of transmission for a pathogen, while transmissibility represents the epidemiological characteristics (e.g. infectious duration, infection probability) that determine effective transmission upon a relevant contact. Consequently, host behavior can affect *R*_0_ which could drive the evolution of pathogen traits.

The associations between contact network structure and pathogen traits are well-supported by theory. For example, sexually transmitted pathogens such as gonorrhea (*Neisseria gonorrhoeae*) or herpes simplex virus rely on rare, dyadic transmission events, likely producing a sparse contact network; to compensate for this sparseness, they are thought to exhibit longer duration infections and higher infection probability respectively (1,5,23). In contrast, tick-borne flaviviruses are only infectious for about 2–3 days in mammal hosts, but persist in tick populations due to their host’s aggregated co-feeding behaviors and consequently high rates of contact (24). Despite these kinds of anecdotal observations, there is no meta-analytic evidence to demonstrate the relationship between transmission routes and pathogen characteristics.

Thus, a critical gap in disease ecology is our understanding of how contact events required for pathogen transmission routes might exhibit distinct contact network structure, and how they might alter the evolution of adaptive pathogen characteristics required to capitalize on these host networks. To answer this question, we conducted a quantitative analysis on 232 animal contact networks spanning eight taxonomic classes to investigate the impact of contact type on pathogen traits. First, we categorized networks into four different horizontal transmission modes based on their contact events. Next, we used a multivariate generalized linear mixed model (GLMM) to identify how network structure is predicted by its associated transmission mode. We then mathematically examined how pathogen traits (i.e. critical transmissibility) may change in order to persist on these different contact networks and compare our results to current knowledge of pathogen traits. We provide practical evidence that contact network structure is influenced by contact events, and that this structural variation causes differences in pathogen transmissibility thresholds that are reflective of our current knowledge of pathogen infection characteristics.

## 2 Methods

We sought to answer the following:

1. How does non-human contact network structure differ depending on the transmission mode associated with its contact event?
2. How does the resulting contact network structure affect a pathogen’s ability to persist on that network?
3. How might these results be reflected in known pathogen traits?

To answer question one, we used a GLMM to examine how contact events associated with different pathogen transmission modes predict eight different descriptors of network structure. To answer question two, we calculated a pathogen’s critical transmissibility (*T*_*c*_) value on these different network types, or the value of transmissibility (*T*) necessary for a pathogen to persist on a network (basic reproduction number (*R*_0_) > 1) where epidemics might occur. To answer question three, we collate published information on pathogen traits in humans that make up *T*_*c*_ (e.g. probability of infection, infectious period) and provide a preliminary comparison between these human pathogen traits, their transmission routes, and our model predictions. Therefore, we aimed to provide evidence that transmission mode affects emergent contact networks, and therefore requires specific evolved pathogen traits to capitalize on.

### 2.1 Dataset

We compiled a dataset of animal contact networks where edges represent one of twelve different contact events, using the Animal Social Network Repository (ASNR) (25,26). The ASNR is an open-source animal behavior network library in which we have compiled network data from the available literature across eight animal taxonomic classes (Mammalia, Aves, Reptilia, Amphibia, Insecta, Arachnida Actinopterygii, and Cephalapoda). Contact events include group membership, nonphysical social interactions, spatial proximity, foraging interactions, trophallaxis (mouth-to-mouth food sharing), synchronous and asynchronous resource sharing, agonistic behaviors, grooming, other physical contact, and mating interactions. Our sample size for this study consisted of 232 contact networks from all eight taxonomic classes (Figure S1). Of these 232 networks, 181 had weighted edges determined by the duration, frequency, or association probability (e.g. half-weight index) of the contact event.

### 2.2 Defining and characterizing contact networks

For each network in our dataset, nodes represented an individual animal and edges represented a contact event between two animals. Based on the contact event, we assigned a transmission mode to each network in our dataset (Table 1). We focus on four transmission modes that our sample represents well: fluid-exchange, direct physical, nonphysical close, and indirect. We define transmission mode as follows:

1. Fluid-exchange: Contact events are behaviors that result in the exchange of bodily fluids. This includes sexual contact such as cloacal transfer, intromission and spermataphore transfer, as well as direct food sharing interactions such as trophallaxis.
2. Direct Physical: Contact events are interactions of physical touch that include grooming, agonistic behaviors (e.g. head-butting, fighting), and other physical social contact.
3. Nonphysical Close: Contact events are in close spatial proximity in which face-to-face contact, or respiratory droplet exchange, could occur. This includes group memberships, or spatial proximity.
4. Indirect: Contact events are asynchronous resource sharing interactions. Indirect contact is unique in that individuals do not need to be using the resource at the same time to be connected in the network.

**Table 1:**
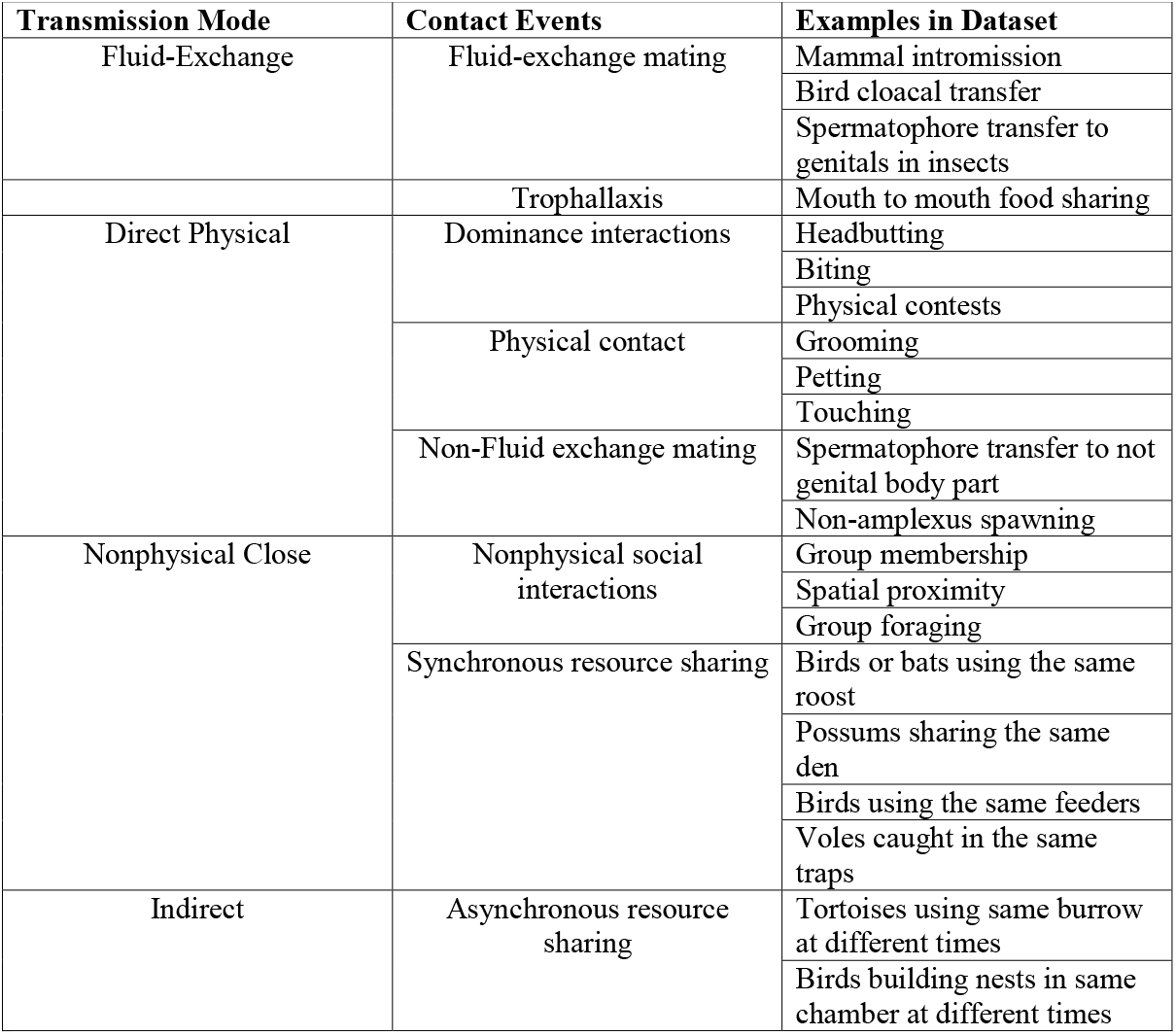
The four transmission modes used to define our 232 contact networks based on the contact event that defined their edges, with examples found in our dataset.

We note that our nonphysical, physical, and fluid-exchange categories have an inherent nested structure with the broadest category being nonphysical, the most selective being fluid exchange (Figure S2). In other words, pathogens that transmit on nonphysical networks (i.e. respiratory droplet pathogens such as SARS-CoV-2) can also transmit on direct physical and fluid exchange networks, but fluid-exchange transmitted pathogens (e.g. sexually transmitted pathogens such as gonorrhea (*Neisseria gonorrhoeae*)) can only transmit on fluid-exchange networks. We manage this nested structure by classifying each empirical network into the most specific category possible using the definitions above.

For each network, we calculated the following eight network metrics that are known to influence infection dynamics and social structure, ignoring edge weights (Table 2): total network density, degree heterogeneity, degree assortativity, average clustering coefficient, average betweenness centrality, network diameter, fragmentation, and subgroup cohesion. Fragmentation (i.e. the number of communities in each network), was estimated using the Louvain method (27) and the remaining network metrics were calculated using the NetworkX package in Python (https://networkx.github.io/).

**Table 2:**
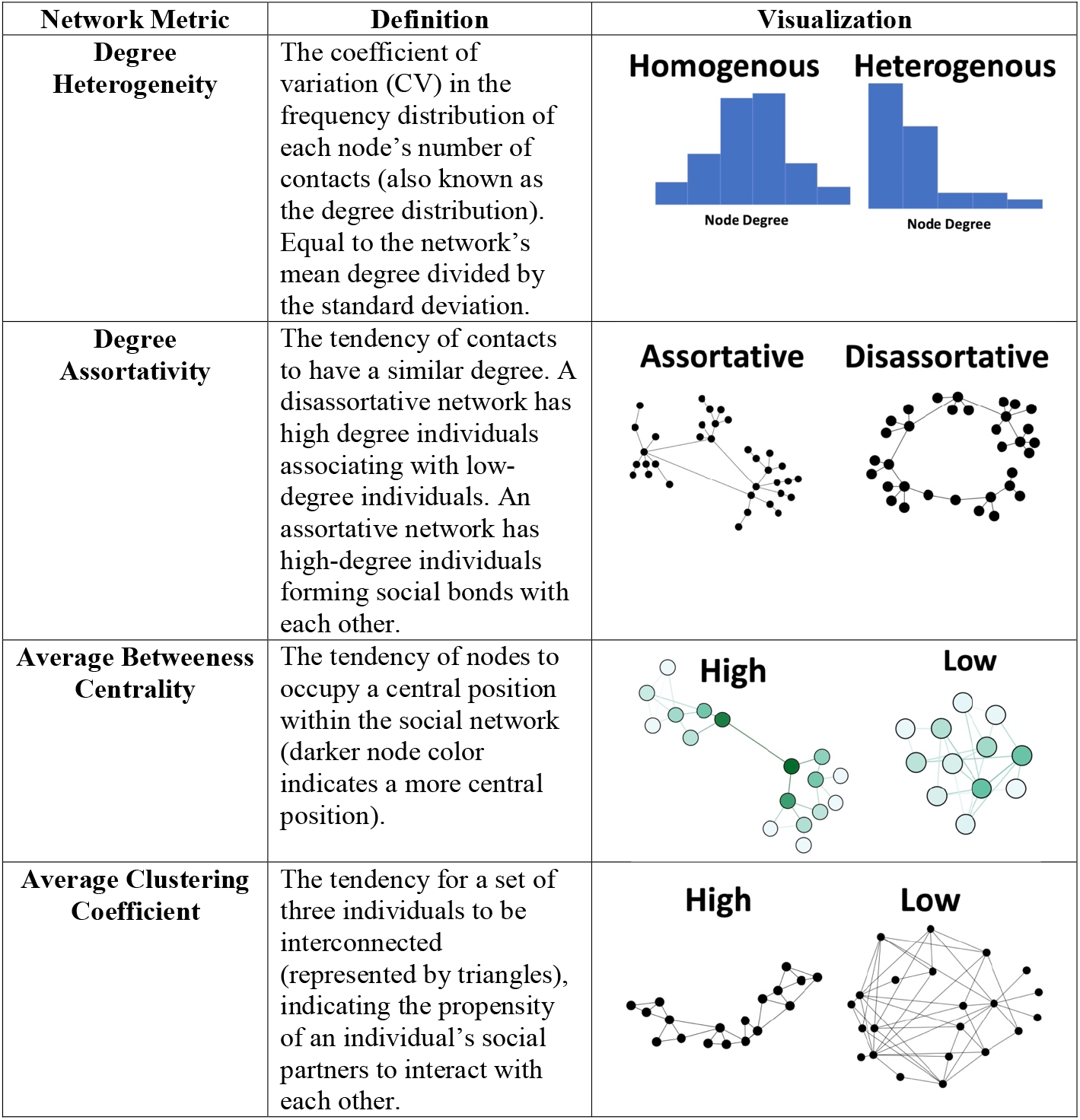
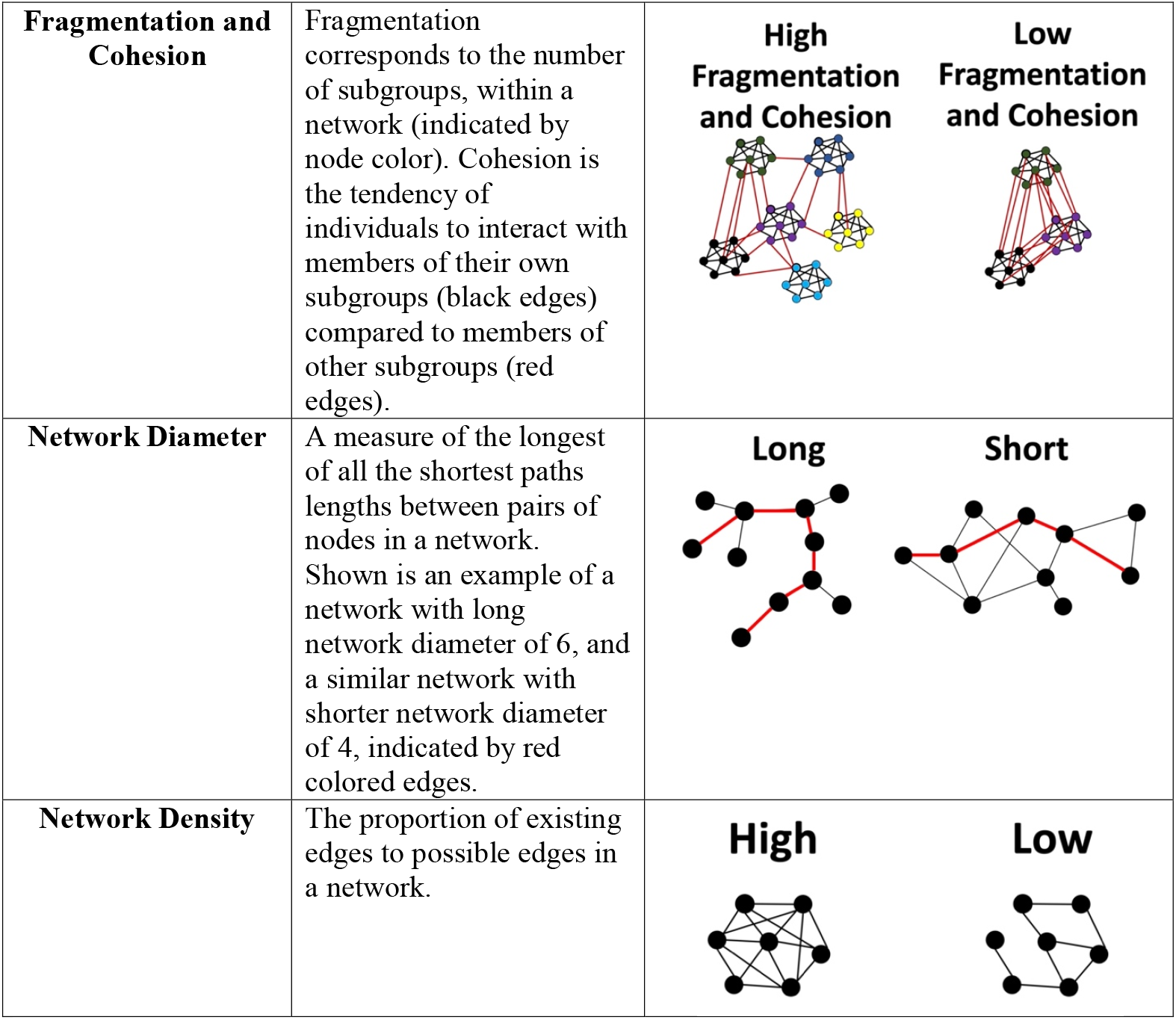
The eight network metric response variables used in the multivariate GLMM: Degree heterogeneity, degree assortativity, average betweenness centrality, average clustering coefficient, fragmentation, cohesion, and network diameter.

### 2.3 Identifying how network structure depends on transmission mode

To examine how contact network structure differs depending on its associated transmission mode, we fitted a multivariate generalized linear mixed model using the MCMCglmm package in R (28), where the eight network metrics (Table 2) made up our multivariate response, and the associated transmission mode was our predictor variable.

We also controlled for the effect of network size on these metrics by including the number of nodes as a predictor. Edge weight type (weighted vs unweighted) was also included to control for data sampling design and edge weighting criteria. As the spatial scale of data collection has been shown to influence network structure (18), we also included sampling scale as a predictor. Studies that collected data on captive animal populations (where all nodes and interactions are theoretically known) were labeled as captive sampling. Studies that focused data collection on specific social groups were categorized as social sampling, and those that focused data collection on all individuals within a fixed spatial boundary were labelled as spatial sampling. Since the social system of an animal species is also shown to influence network structure (18), we included species social structure (relatively solitary, gregarious, and socially hierarchical defined in Table S1 based on (18)) as a predictor. Finally, we controlled for repeated measurements within studies by including study id as a random effect in the analysis. We were unable to include a random effect for taxonomic class or use a phylogeneticaly controlled model, because there was an unbalanced representation of different taxa across the four transmission mode categories that made these effects difficult to fit successfully (Figure S1).

All response variables were continuous; to encourage proper model fitting we log-transformed then centered them (by subtracting the mean) and then scaled to unit variances (by dividing by the standard deviation). We ran 1 MCMC chain for 10,500 iterations, with a thinning interval of 10 after burn-in of 500 with uninformative priors. Non-physical contact transmission was the intercept factor level for the transmission mode fixed effect. For each response variable, if the effect sizes of the three remaining transmission modes overlapped with zero then it was considered not different from non-physical contact. To examine differences between the remaining three modes, we observed the proportional overlap between their effect sizes across all 1000 iterations, multiplied by 2 per a two-tailed test; if it was less than 0.05, then the response variables were considered different between the transmission modes.

Within the ASNR, several studies provided more than one network for the study species or population. To avoid biasing our sample towards these studies, we randomly selected a maximum of 15 networks from each study (n=232). To ensure that our results were not affected by this random subsampling, we reran our model 1000 times, each time choosing a different sample of networks from each study with more than fifteen networks. We then chose a random estimate from each model run and computed an average of model estimates across the 1000 different subsamples.

#### 2.3.1 Investigating the role of weak ties

To determine if weak ties (i.e. low edge weights) might drive contact network structure across transmission modes, we recalculated all eight network metrics for each network in our dataset when the lowest 5, 10 and 15% of weighted edges were dropped from the network. We then reran our GLMM to see if differences in network metrics among our different transmission modes still hold true when low weight edges are no longer accounted for.

### 2.4 Characterizing critical transmission thresholds

To examine how network structure affects a pathogen’s ability to persist on a network, we sought to identify a pathogen’s critical transmissibility (*T*_*c*_) on contact networks based on transmission mode. For a pathogen to persist in a network, its basic reproduction number (*R*_0_) must be greater than 1, and *R*_0_ depends on both a pathogen’s *T* and the contact patterns of the network it travels on. Therefore, for each network, we sought *T*_*c*_ for which *T*_*c*_ *(contact)* = *R*_0_ > 1. To estimate *T*_*c*_, we considered the impact of contact network structure by calculating *R*_0_ using Monte Carlo simulations of a susceptible-infected-recovered (SIR) model of infection spread through each network. We ignored edge weights because the impact of interaction weight (e.g. contact duration or frequency) on infection spread is not well understood generally. We used an SIR percolation simulation model (29), where each outbreak was initiated by infecting a randomly chosen individual in the network. For the first generation of the simulation, the individual is given an opportunity to infect all its contacts, with transmissibility *T*, and then recover. This process is then repeated for each infected node, until no infections remain in the network. For each network, we simulated 250 disease outbreaks and of those classified large-scale epidemics as those where at least 10% of the population is infected. We repeated this for each *T* value in the range (0.01-0.8), and recorded the first *T* value for which at least 10% of the outbreaks were largescale epidemics. (Percolation theory suggests that our expectation for the probability of having a largescale epidemic should match the expected size of a large-scale epidemic (29)). This reported *T* value is our estimate of the network’s critical transmissibility *T*_*c*_.

Past studies suggest that degree and degree heterogeneity are the most important aspects of network structure affecting how a pathogen will transmit across a network (30). In order to test this, we considered two control scenarios where we 1) isolated the effect of homogeneous degree (i.e. all individuals have the same number of contacts) on *T*_*c*_ and 2) isolated the effect of heterogeneous degree (i.e. on average, individuals have the number of contacts as scenario 1, but individual degree varies around this mean) on *T*_*c*_. Therefore, we sought *T*_*c*_ for which *T*_*c*_ ⟨*k*_*e*_⟩ = *R*_0_ > 1, where ⟨*k*_*e*_⟩ is the average excess degree of the network. The excess degree is the potential number of contacts an individual can infect after they have been infected by one of their contacts, and on average this value is larger for networks with degree heterogeneity than for homogeneous degree networks (22).

For the first control scenario, we considered a pathogen’s *T*_*c*_ in a homogeneous degree network, in which the average excess degree, ⟨*k*_*e*_⟩, is:

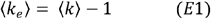

where ⟨*k*⟩ is the average degree of the network. For the second control scenario, we considered a network with degree heterogeneity, thus the average excess degree is:

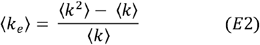

We compared the *T*_*c*_ values for each transmission category of contact networks (non-physical close, direct physical, fluid-exchange, indirect) within each scenario using a one-way ANOVA, and pairwise t-tests with a Tukey HSD familywise error-rate correction.

### 2.5 Examining diversity of empirical pathogen characteristics

To examine how our results are reflected in known pathogen traits, we considered the transmissibility of a pathogen (*T*) as a function of its infectious duration (*G*) and its probability of infection (*β*) (21,22):

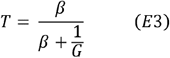

To provide context for the covarying characteristics of known pathogens, we examine known β and G values (infection characteristics) for a small set of well characterized pathogens to see how they compare to our findings. Because data on pathogen traits in non-human animals is limited, we instead focus on these traits in common human pathogens. We use two systematic reviews on the natural histories of pathogens for common diseases in preschools (31,32) to summarize data on pathogens that use each transmission mode. We only included the pathogens in our summary if the source of the data was from a well-designed study, using the levels of evidence I and II provided in the first review (31) (Table S2). If there was no data, or the source of the data was poor (level of evidence III or IV) for the pathogen’s infectious period (*G*), we instead used the shedding periods, defined as the period of time during which an individual excretes the pathogen; the shedding period can be used to estimate the duration of infectiousness when there is lack of direct evidence (31). If there was no or poor data for the shedding period, then it was not included in this summary. We then verified the data for these pathogens using the second review (32).

Since these reviews did not contain any sexually transmitted pathogens, we took the ten sexually transmitted pathogens listed on the CDC website (www.cdc.gov/std) and examined the literature for their natural histories. We found studies that estimated both the probability of infection (*β*) and infectious period (*G*) of four of the ten pathogens (syphilis (33,34), gonorrhea (35,36), chlamydia (37,38), and trichomoniasis (39,40)).

## 3 Results

### 3.1 Network structure is dependent on behavior type

We examined how network structure was predicted by the pathogen transmission mode represented by specific contact events using a multivariate GLMM. We summarized network structure with eight topological characteristics (Table 2), and all the metrics except subgroup cohesion and fragmentation differed among transmission mode categories. The predicted distributions for the six remaining network metrics by transmission category are summarized in Figure 1 and all other effect sizes from our model are in Tables S3, S4, S5, and S6.

**Figure 1:**
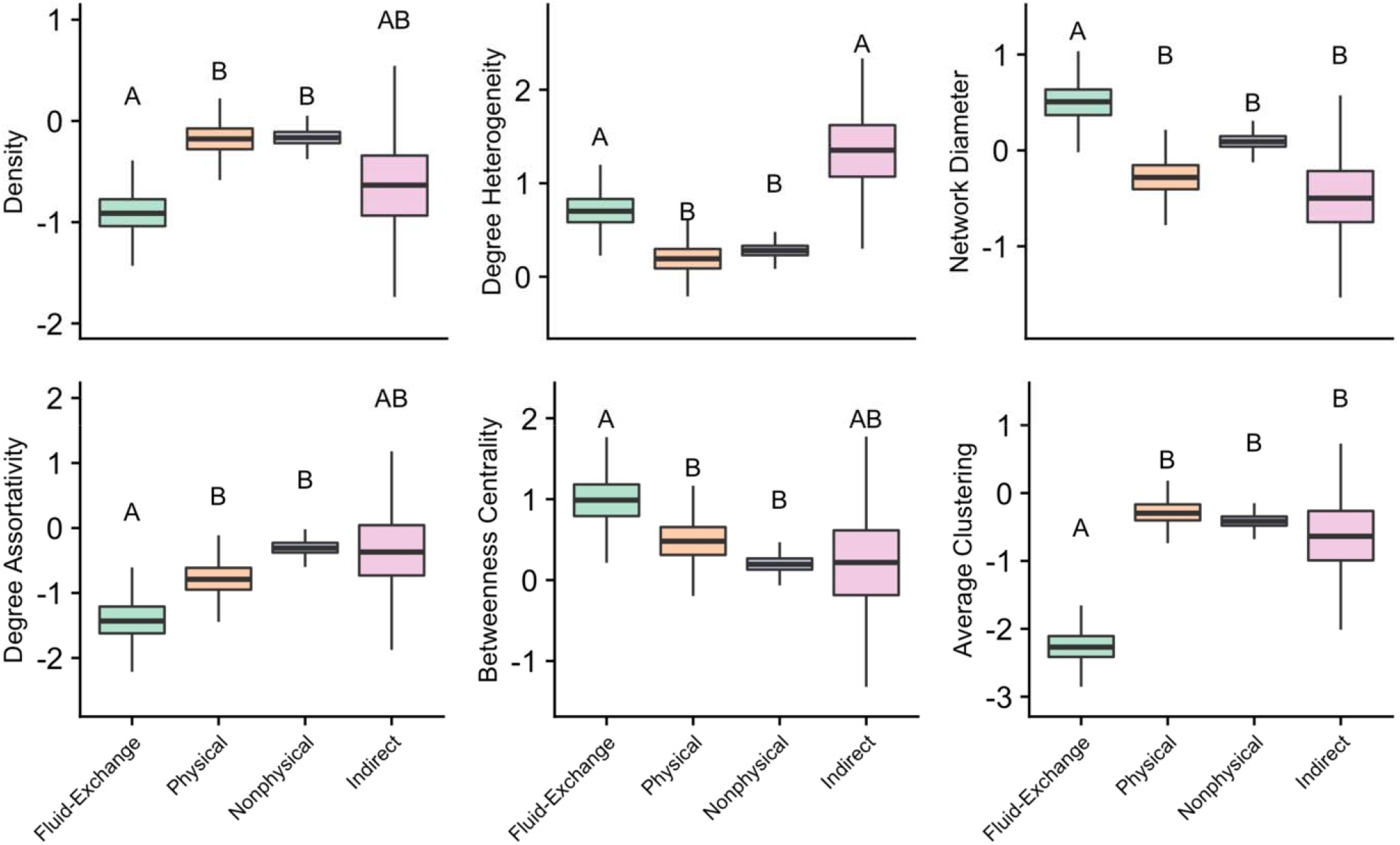
The predicted distributions of six of the eight network metrics for each transmission mode, based on the results of the GLMM. Letters represent significant differences between transmission modes. Results for fragmentation and cohesion were not different among transmission modes and are therefore not included in this figure.

Fluid-exchange contact networks differed from physical and nonphysical contact networks in all six network metrics and differed from indirect contact networks for clustering and network diameter. Indirect contact networks only differed from physical and nonphysical networks in their degree heterogeneity. Physical and non-physical networks could not be differentiated by the network metrics we tested. This suggests that fluid exchange contact events create the most unique contact patterns.

Physical and nonphysical contact networks had higher density values and shorter network diameters than fluid-exchange networks, and lower degree heterogeneity than both fluid-exchange and indirect contact networks. This indicates that physical and nonphysical contact events create networks that are more connected, with less variation in each individual’s number of contacts.

Fluid-exchange contact networks also had lower degree assortativity than physical and nonphysical contact networks. That is, the contacts of high degree individuals in fluid-exchange contact networks were more likely to be low degree than in physical and nonphysical contact networks. Indirect contact networks did not differ in their degree assortativity from any other contact network. Fluid-exchange contact networks also had lower average clustering values than all other contact networks.

Finally, fluid-exchange contact networks have higher betweenness centrality values than physical and nonphysical contact networks. This means that these networks tend to have more “bridge nodes”, or nodes that connect different communities together: this is despite the fact that the number of communities (fragmentation) and the cohesiveness of those communities do not differ among contact networks.

#### 3.1.1 Weak Ties

To examine the role of weak interactions (i.e. low edge weights) in determining unique network structure, we reran the GLMM on our dataset after dropping the lowest 5%, 10%, or 15% of weighted edges in each network. We found that the significant differences in average clustering and degree assortativity in networks across the four transmission categories are maintained as we filter low weight edges. However, as edges are filtered, there are no longer any differences in density and degree heterogeneity (Figure S3) among contact network categories. In other words, without low weight edges, contact networks in each transmission category become more similar in their number and variation of contacts.

### 3.2 Pathogen transmissibility must be higher for contact networks with lower connectivity

We demonstrated the effect of contact on critical transmissibility values (*T*_*c*_) in each of our four contact network categories by considering their full unique network structure. We then specifically examined the effect of network density (average number of contacts) and degree heterogeneity (variation in number of contacts) on *T*_*c*_ in two different control scenarios.

We find that generally pathogens needed significantly higher critical transmissibility values on fluid-exchange contact networks than on physical, nonphysical, and indirect transmission contact networks (Figure 2). By comparing our empirical simulation scenario to the two control scenarios, we find that fluid exchange and indirect contact networks are more vulnerable to disease invasion (i.e. lower *T*_*c*_) than expected based on their average connectivity (Figure 2). This result is consistent with network epidemiology theory which predicts that higher degree heterogeneity (as we find in fluid-exchange and indirect contact networks) make disease invasion more likely. For physical and non-physical contact networks, on the other hand, the critical transmissibility is comparable in all three scenarios, suggesting that the average network connectivity (or network density) is sufficient to predict disease invasion in such networks.

**Figure 2:**
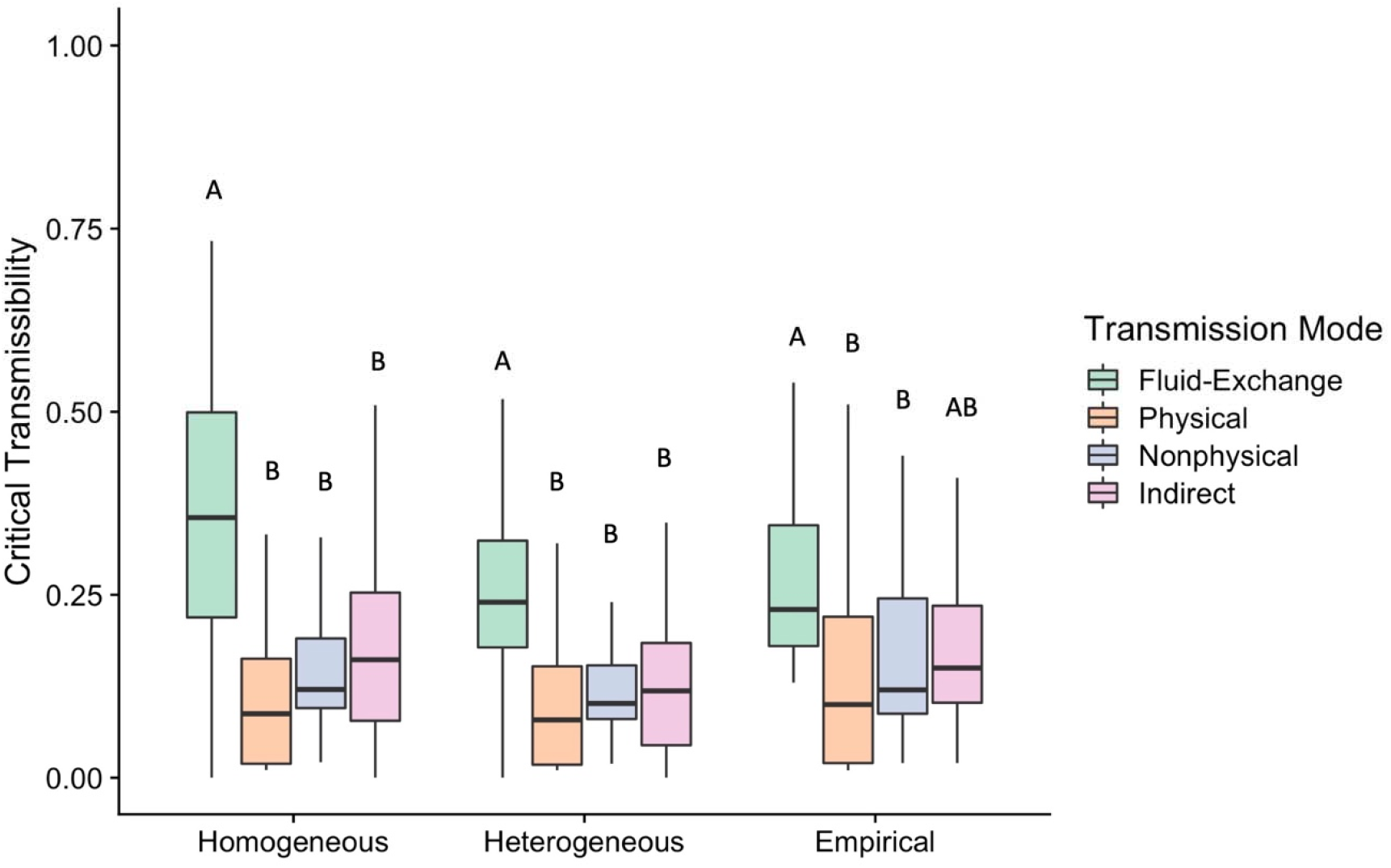
The critical transmissibility values (*T*_*c*_) needed for a pathogen to persist (*R*_0_ > 1) on each contact network category. *T*_*c*_ was estimated empirically using SIR simulations on each network to account for the effect of full network structure. Tc was then calculated in two control scenarios that examined the effects of homogeneous degree (E1) and heterogeneous degree (E2). Colors represent the different contact network categories, and letters indicate significant differences within each method based on a One-Way ANOVA and Tukey’s HSD

### 3.3 The structure of a contact network can influence the infection characteristics of associated pathogens

We compiled peer-reviewed data from common human pathogens as life history data on pathogens in non-human animals was extremely limited. We visualized two of the characteristics that define a pathogen’s transmissibility: its infectious period and its infection probability (Figure 3). First, we showed that pathogens that transmit on indirect contact networks (e.g. food/waterborne, fecal-oral) have relatively short infectious periods, and varied between low to moderate infection probability. Next, we found pathogens that transmit on fluid-exchange contact networks (e.g. sex, saliva) have the longest infectious periods. Of those, the two pathogens that tend to have shorter infectious periods alternatively have moderate to high infectivity (herpes, syphilis). These results suggest that fluid-exchange pathogens do in fact have higher *T*_*c*_ values than other pathogens, and they tend to increase their *T*_*c*_ by extending their infectious periods over increasing their infection probability. Finally, we see that pathogens that transmit on physical and nonphysical contact networks have some of the shortest infectious periods and a range of infection probabilities, but most pathogens with high infectivity are associated with these transmission modes.

**Figure 3:**
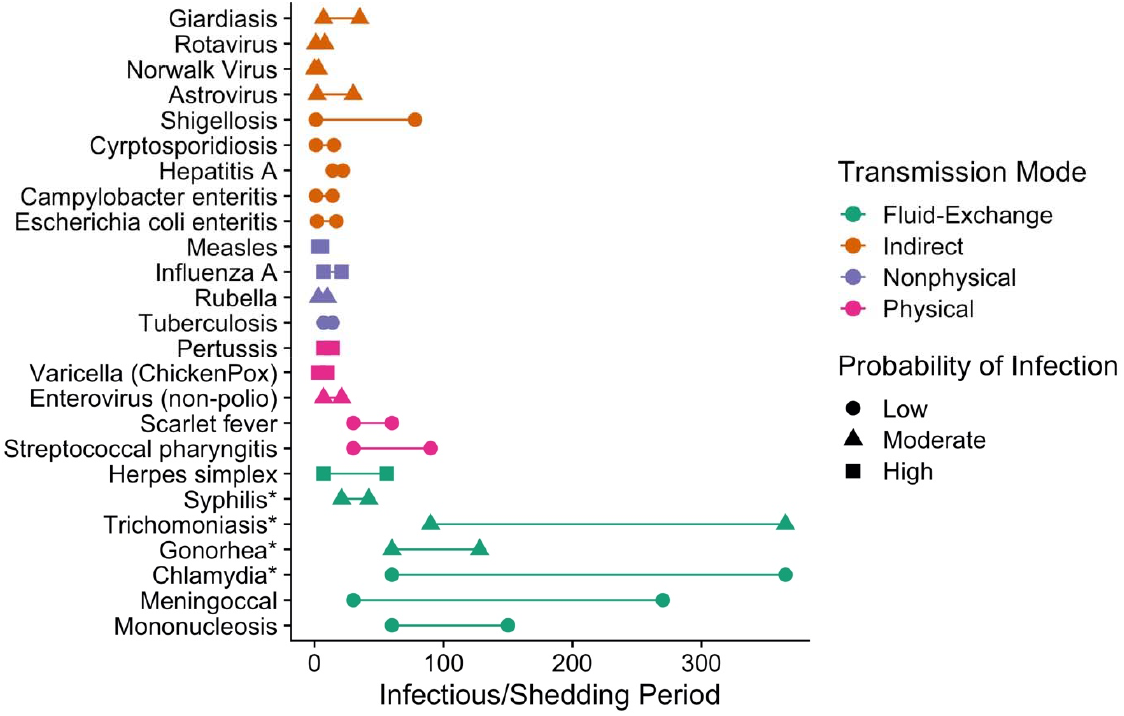
The typical infectious periods of 25 different human pathogens from two literature reviews (31,32) (pathogens with asterisks came from alternative sources). Color denotes the transmission mode of each pathogen. Shapes indicate the pathogen’s probability of infecting a host given contact.

## 4 Discussion

Our study shows that networks characteristic of different pathogen transmission modes differ in terms of their structure. We also show that differences in network structure will affect the transmissibility required for a pathogen to successfully proliferate. Finally, we suggest these network structures likely impact the evolution of a focal pathogen’s infection characteristics, supported by a review of human pathogen traits.

### 4.1 Differences among contact networks and their implications for pathogen spread

Most notably, compared to physical and nonphysical contact networks, fluid exchange networks were less dense and had more heterogeneous degree values, with greater diameters, reduced clustering, and greater disassortativity by degree. They also had more bridge nodes than all other network types. That is, these networks tend to be more poorly connected, and with greater skew in nodes’ importance in the network. This selection of network traits likely arises through a combination of mechanisms, linked to the fact that many of these networks were based on sexual interactions.

First, the infrequency of copulation events will drive low network density, with correlated increases in the diameter of the network and the prevalence of bridge connections; second, the fact that all networks solely included male-female copulation events likely increase clustering and reduce the tendency for assortative mating; and finally, the common nature of polygyny and overdispersed mating events (i.e., few individuals monopolizing sexual resources) will drive greater degree heterogeneity, as well as driving disassortativity. These latter traits agree with our understanding of the overdispersed nature of sexual interaction networks (41–43). The fluid-exchange category also included trophallaxis networks, which are also generally highly heterogeneous; in ants, more than 50% of trophallaxis interactions may come from less than 25% of individuals (44). While the majority of the trophallaxis networks represented in our dataset are indeed in ants, many other species such as birds and mammals also partake in trophallaxis, usually in the form of parental care (45); these unique parent/offspring interactions would also likely result in sparse, heterogeneous, and highly fragmented networks.

These traits could have a selection of important consequences for transmission of pathogens through fluid exchange networks. First, less dense networks are likely to provide fewer transmission opportunities for fluid-borne pathogens (46), while larger diameters will inhibit the spread of an outbreak. Second, degree heterogeneity is known to be key driver of sexually transmitted pathogen risk in human contact networks (16,47). Where contact numbers are highly heterogeneous, superspreaders can lead to rapid, explosive outbreaks, allowing pathogens to persist (21). Indeed, past work has shown that HIV, a sexually transmitted pathogen, can exploit this contact heterogeneity to attain sufficient transmissibility, and other sexually transmitted pathogens likely do the same (48). However, previous work on contact networks has shown that network density and individual variation in contact are negatively correlated (49). Our results support these findings as fluid-exchange and indirect contact networks are shown to have lower densities and higher degree heterogeneity compared to physical and nonphysical contact networks. This suggests that while having high degree heterogeneity might make a network more vulnerable to explosive outbreaks (18,21), this may trade off with lower overall transmission probabilities (49). Third, low clustering values might be beneficial for pathogens, especially given the low densities, as a pathogen may be less likely to get stuck in cliques that might form among individuals (50). Fourth, the common nature of bridge nodes in fluid exchange networks might be especially important when considering control measures for pathogens on these networks. Previous work has shown that pathogens with high transmissibility are able persist in socially fragmented networks because bridge nodes allow for transmission amongst communities (30). Therefore, the removal of these nodes will prevent or slow pathogen spread through a population (47). While both physical and fluid exchange transmission require relatively close contact, controlling disease spread by identifying bridge nodes might be more powerful for fluid-exchange pathogens than for others.

Notably, indirect contact networks had a more heterogeneous degree distribution than physical and nonphysical contact networks, meaning there is more between-individual variation in asynchronous resource sharing compared to that in close proximity or physical touch. Past research has demonstrated this phenomenon in many solitary desert species that asynchronously burrow share, and while there are many ecological factors that might drive these individual preferences for resource use patterns (such as differences in sex, age, and environment) (30,51), we still do not have a full mechanistic understanding of them. It is possible that this pattern will drive greater heterogeneity in infection with indirectly transmitted pathogens.

### 4.2 Network structure and the pressures on pathogen characteristics

Pathogens that transmit on indirect contact networks (e.g. food/waterborne, fecal-oral) seem to have relatively short infectious periods, and vary between low to moderate infection probabilities (Figure 3). However, we found many aspects of indirect contact network structure were not different from fluid-exchange networks (i.e. low network densities and high degree heterogeneity) (Figure 1). These results contradict expectations that these indirect networks should be highly connected since individuals need to have only used the same space at some point in time to be connected. For example, many individuals sharing the same sanitation facilities through time such as on airplanes (52), cruise ships (53), and hotels (54) can cause recurring outbreaks of Norwalk virus, a common fecal-oral pathogen; since there is no need to physically contact an individual to be infected by them, these contact networks are known to be extremely highly connected (55). There are several possible explanations for our surprising finding: first, our results may be driven by the fact that relatively solitary species (which have low connectivity due to their social structure (30)) are highly represented in our indirect networks sample. Additionally, these networks often involve territorial species; resource sharing (both synchronous and asynchronous) is likely minimized in species that hold territories (e.g. (56)). Moreover, territoriality can be sex specific in that males very rarely use the same space and resources even asynchronously but females can move freely between male territories, which can result in a sex-specific degree heterogeneity in some species (e.g. (57,58)). Indeed, our GLMM showed a high amount of variation in the effect sizes of indirect contact networks and we found that pathogens using indirect contact networks do not need high *T*_*c*_ values to persist on these networks. This suggests that a species’ social system strongly influences the structure of indirect contact networks. Including additional species with other social systems may increase the average connectivity of indirect contact networks, which would be more representative of their associated pathogen characteristics. This paucity of variation in social systems is a common problem in meta-analyses of social network structure, and ongoing data collection may help to ameliorate this difficulty in the future.

Our simulations revealed that contact network structure should motivate the evolution of higher transmissibility for fluid exchange pathogens to persist (Figure 2); a prediction that was supported by our literature review (Figure 3). This supports what we know about the behaviors involved in bodily fluid exchange; individuals usually have long temporal gaps in between fluid exchange events compared to other potential disease transmission behaviors (1). Therefore, pathogens would benefit more from longer infectious periods giving them more time to spread.

### 4.3 The role of weak ties in defining relevant disease spreading contact

Previous studies have shown that structural differences between networks are primarily driven by “weak ties” that are disproportionately lower in intensity, frequency, or duration than other contacts (18). When we eliminated weak ties, we found that differences between transmission types persisted for some structural features (e.g. average clustering and degree assortativity), but others (density and degree heterogeneity) were lost; in other words, removing the weak ties from contact networks makes them more similar to each other in their number and variation of contacts. Given that the structural features of density and degree heterogeneity were most different among fluid-exchange networks, this finding suggests that not only do individuals tend to vary in the number of fluid-exchange contacts they have compared to their other types of contacts, but individuals also vary more in the strength of connections between their different fluid-exchange contacts, compared to their other types of contacts. We hypothesize that this might be the crucial difference between pathogens that spread via fluid exchange and others, but additional data would be required to confirm this hypothesis.

This heterogeneity in how individuals distribute their contact effort is known as “social fluidity”, where higher social fluidity suggests a higher presence of weak ties (9). Past work has shown that nonphysical contact networks (e.g. spatial association) have smaller values of social fluidity than fluid-exchange networks (e.g. trophallaxis), suggesting that social fluidity and weak ties are especially relevant when considering disease transmission potential in these networks (9). In some instances, not considering a very brief or infrequent contact as “relevant” for disease transmission might makes sense, such as for pathogens that propagate on networks with low social fluidity (e.g. flu). However, for pathogens spread via fluid exchange, we found that that the density and degree heterogeneity of contact networks are important predictors for determining their transmissibility and traits. This suggests it is imperative to include short or infrequent fluid exchange interactions when considering the definition of relevant contact for modeling the transmission of these pathogens. By not including these “weak” contacts, it is likely that estimates of fluid-exchange pathogen spread would be inaccurate, and proposed control measures based on these estimates could be unreliable.

### 4.4 Study Limitations

Our study has some important limitations. First, we investigated non-human animal networks, while using pathogen traits in humans to understand the implications of network structure on infection characteristics. We would expect some aspects of human contact networks to differ from animal contact networks, particularly with fluid-exchange transmission. For example, human sexual networks have much higher clustering values than what we observed in our animal networks (59). However, this could be due to the lack of recorded same-sex sexual behaviors in non-human animal species; same sex behaviors including fluid exchange do occur in non-human animals and are therefore likely underrepresented in our sample (60). Regardless, even if true clustering values are higher than observed, we would expect this to *reduce* the *R*_0_ of a pathogen in a network; this might further increase the transmissibility needed to persist (59), further supporting our current results. Overall, we found that network density and degree heterogeneity were the most important metrics when considering a pathogen’s ability to persist on a network; we show that fluid-exchange contact networks have low densities and high degree of variation which holds true in human networks (1,16,47). Future studies could further validate this work by exploring the contact structures of available human contact networks. Alternatively, future work could provide a better overview of the natural histories of pathogens in wildlife. While we would expect infection characteristics to be similar based on our network analysis, we suggest a more thorough review of these characteristics across different taxa.

Our available network datasets likewise restricted our ability to test and untangle some factors. First, because we had a lack of data on indirect contact networks in wildlife species, we were unable to investigate the evolution of traits of environmentally transmitted pathogens. We also did not have good representation of contact networks representative of each transmission mode across all taxonomic classes, which reduced our ability to control for host taxonomy in our model. Future work may be able to ameliorate these difficulties by measuring indirect contact networks from species with gregarious and hierarchical social makeups, as well increasing the taxonomic sample size of each transmission mode category, for a better representation of contact networks across species and social systems.

Lastly, our work does not consider the impacts of pathogen-mediated changes to contact structure as caused by sickness behaviors due to host immune response or pathogen virulence, nor due to pathogen manipulation of host behavior ((15), and e.g. (61,62)). Such disease-mediated behavior change can alter realized pathogen characteristics (e.g. infectious periods can be effectively reduced via sickness behaviors). Future work must consider this critical feedback loop between contact structure and pathogen characteristics, in light of transmission modes.

## Acknowledgements

We thank Pratha Sah, Janet Mann, and Tommaso Pizzari for their thoughtful comments on this work. We also thank Pratha Sah, Jose Mendez, Grant Rosensteel, Elly Meng, and Sania Ali for their contributions to the development and growth of the Animal Social Network Repository. This work was supported by the National Science Foundation (Award # DEB-2211287) and the Morris Animal Foundation (Award # D22ZO-059).

## Data Availability

All networks used in this study can be found in the Animal Social Network Repository (25). All code can be found at (https://github.com/mac532/pathogen-transmission-contact-network).

