## Supplementary Information for "Pathogen transmission modes determine contact network structure, altering other pathogen characteristics"

Table S1: The definitions for the social system categories used as predictor variables in the GLMM for each contact network. Definitions are from (18).

| Social System | Definition |
|---|---|
| Relatively Solitary | Infrequent aggregation or association between adults outside of the breeding period, and lack of synchronized movements in space by adults |
| Gregarious | Species that aggregate for one or more activities, but have unstable or temporally varying group composition |
| Socially Hierarchical | Species characterized by a permanent or long-term (i.e. at least over a single breeding season) stable social hierarchy |

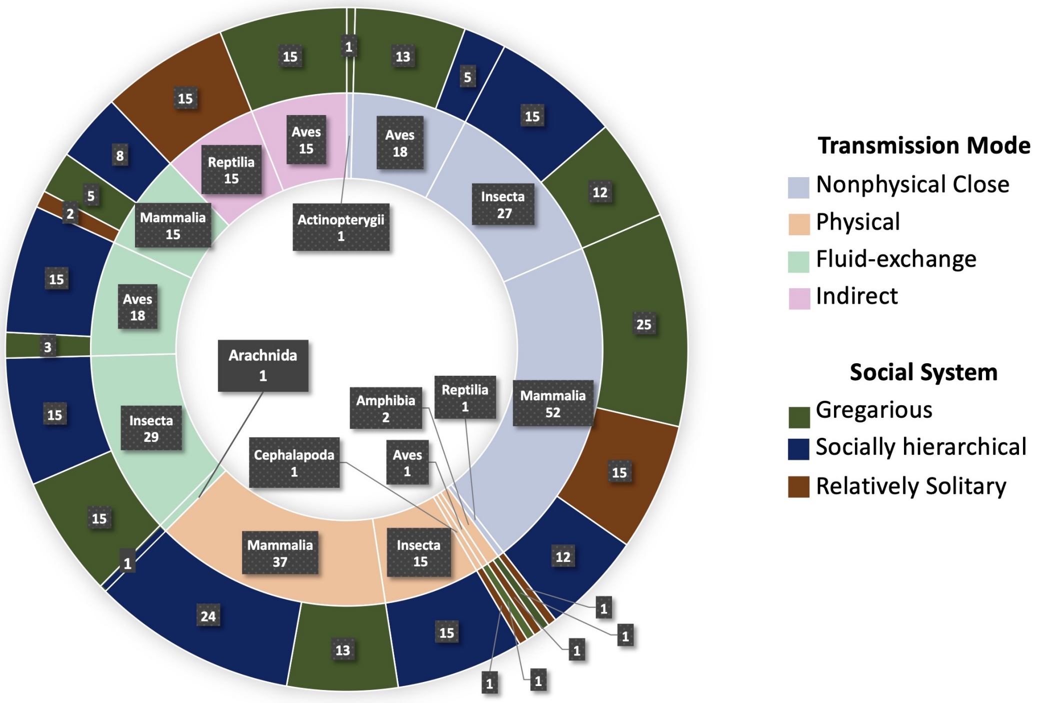


Figure S1: A breakdown of the 232 networks used in our GLMM by taxonomic class and social system. Networks were first categorized into one of four different transmission modes based on the contact event described by the network (Table 1) (inner ring colors). The inner ring also shows the number of networks in each taxonomic class for each transmission mode category. The outer ring shows the number of networks included in each social system, for each taxonomic class.

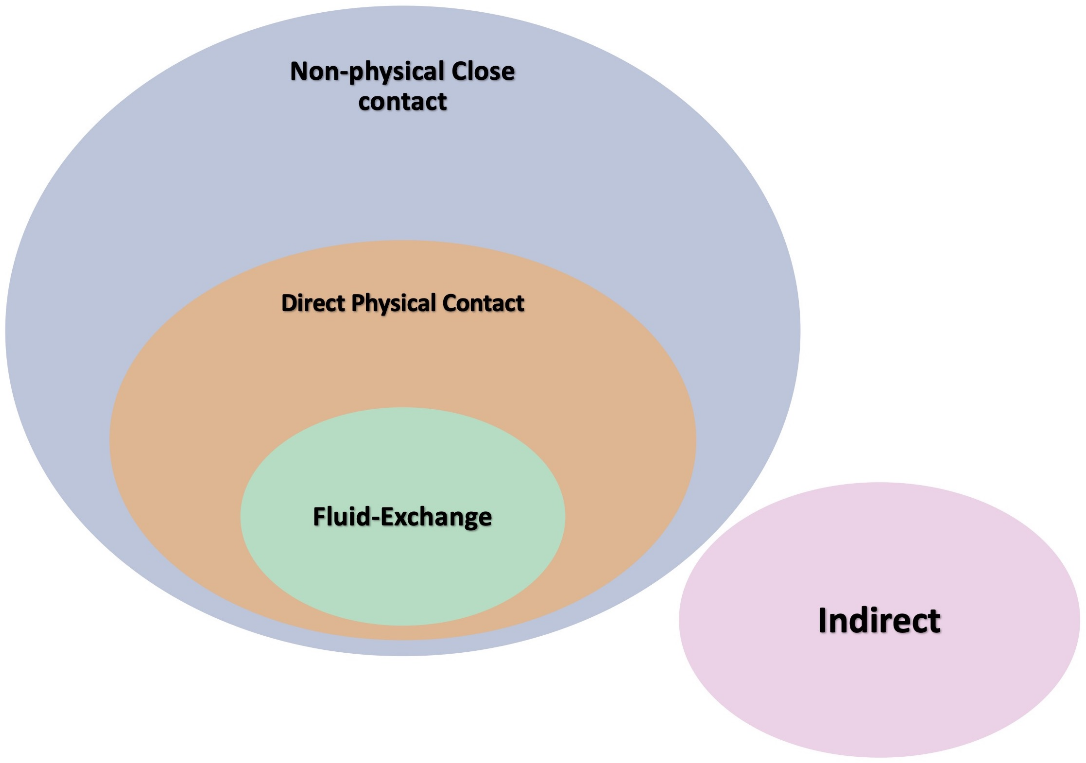


Figure S2: The nested structure of our transmission mode classification. Most networks can be classified in the non-physical close transmission category, the broadest classification on which airborne or respiratory droplet transmitted pathogens can travel. The addition of direct physical transmission is specific to pathogens that transmit via skin-to-skin contact, but airborne or droplet transmitted pathogens can still spread on these networks. Similarly, networks in the fluid exchange transmission category have contact events that involve physical contact with fluid exchange on which sexual fluid or saliva exchanged pathogens can travel, but skin-to-skin and respiratory transmitted pathogens can also still spread on these networks. Indirect contact is outside this nested transmission structure as the contact type is asynchronous (e.g. desert tortoises share burrows, but at different times and will therefore never even be spatially associated).

Table S2: The levels of evidence used in (31) to define pathogen traits. We only included pathogens from which traits came from levels of evidence I and II.

| **Level** | **Definition** |
|---|---|
| I | Systematic review, metaanalysis or well-designed epidemilogic or experimental study with $\geq$ 50 subjects |
| II | Well-designed epidemilogic or experimental study with 5 - 50 subjects |
| III | Case reports with $<$ 5 subjects, or poorly substantiated larger study |
| IV | Opinion or clinical experience of experts (not supported by published data) |

Table S3: Effect size estimates of the generalized linear mixed models continuous effects, network size and duration of data collection, on eight network metrics and the 95% credible intervals included in brackets. For each network metric, if the credible intervals do not overlap zero, the effect was considered significant.

| Network Metric | Variable | Effect Size | 95% credible interval |
|---|---|---|---|
| Density | Duration of data collection | 0.004 | [-0.059, 0.080] |
| | Network size | **-0.306** | **[-0.767, -0.221]** |
| Degree Heterogeneity | Duration of data collection | -0.072 | [-0.141, 0.001] |
| | Network size | **0.183** | **[0.867, 0.467]** |
| Betweeness | Duration of data collection | -0.051 | [-0.156, 0.049] |
| | Network size | -0.036 | [-0.539, 0.049] |
| Degree Assortativity | Duration of data collection | 0.034 | [-0.118, 0.104] |
| | Network size | **0.132** | **[0.012, 0.670]** |
| Diameter | Duration of data collection | 0.007 | [-0.088, 0.075] |
| | Network size | **0.212** | **[0.065, 0.556]** |
| Clustering | Duration of data collection | 0.006 | [-0.080, 0.082] |
| | Network size | -0.043 | [-0.149, 0.101] |
| Cohesion | Duration of data collection | 0.022 | [-0.121, 0.131] |
| | Network size | -0.175 | [-0.596, 0.129] |
| Fragmentation | Duration of data collection | -0.079 | [-0.154, 0.012] |
| | Network size | **0.291** | **[0.140, 0.738]** |

Table S4: Model estimates for the edge weight type co-variate on eight different contact network metrics and their 95% credible intervals in brackets, where "unweighted" was the intercept. For each network metric, if the credible intervals for weighted networks overlap zero, then it was considered not different from unweighted networks.

| Network Metric | Intercept | Focal:Weighted |
|---|---|---|
| Density | Unweighted | **0.390 [0.028, 0.747]** |
| Degree Heterogeneity | Unweighted | -0.157 [-0.512, 0.201] |
| Betweeness | Unweighted | 0.023 [-0.448, 0.679] |
| Degree Assortativity | Unweighted | -0.074 [-0.605, 0.584] |
| Diameter | Unweighted | -0.049 [-0.498, 0.362] |
| Clustering | Unweighted | 0.232 [-0.203, 0.711] |
| Cohesion | Unweighted | **1.136 [0.477, 1.821]** |
| Fragmentation | Unweighted | 0.242 [-0.205, 0.651] |

Table S5: Model estimates for the sampling scale co-variate on eight different contact network metrics with their 95% credible intervals in brackets. This model was run twice, once with "captive sampling" as the intercept and again with "social sampling" as the intercept. For each network metric, if the credible intervals for the focal sampling scale overlap zero, then it was considered not different from the intercept.

| Network Metric | Intercept | Focal:Social | Focal:Spatial |
|---|---|---|---|
| Density | Captive | 0.057 [-0.352,0.352] | **-0.521 [-0.937, -0.214]** |
| | Social | | **-0.620 [-0.862, -0.213]** |
| Degree Heterogeneity | Captive | -0.034[-0.401, 0.320] | **0.336 [0.037, 0.743]**] |
| | Social | | **0.480 [0.144, 0.794]** |
| Betweeness | Captive | 0.474 [-0.194, 0.866] | -0.286 [-0.761, 0.237] |
| | Social | | **-0.599 [-1.027, 0.076]** |
| Degree Assortativity | Captive | 0.199 [-0.419, 0.680] | 0.398 [-0.109, 0.929] |
| | Social | | 0.310 [-0.301, 0.760] |
| Diameter | Captive | 0.002 [-0.391, 0.364] | 0.029 [-0.329, 0.480] |
| | Social | | 0.149 [-0.302, 0.442] |
| Clustering | Captive | 0.138 [-0.384, 0.586] | -0.309 [-0.749, 0.175] |
| | Social | | **-0.533 [-0.867, -0.0573]** |
| Cohesion | Captive | 0.055 [-0.555, 0.609] | -0.553 [-1.079, 0.040] |
| | Social | | **-0.499 [-1.112, -0.032]** |
| Fragmentation | Captive | -0.040 [-0.378, 0.586] | -0.098 [-0.637, 0.182] |
| | Social | | -0.180 [-0.544, 0.191] |

Table S6: Model estimates for the social system co-variate on eight different contact network metrics with their 95% credible intervals in brackets. This model was run twice, once with "Relatively solitary" as the intercept and again with "Gregarious" as the intercept. For each network metric, if the credible intervals for the focal system overlap zero, then it was considered not different from the intercept.

| Network Metric | Intercept | Focal:Gregarious | Focal:Socially Hierarchical |
|---|---|---|---|
| Density | Relatively Solitary | 0.246 [-0.344, 1.003] | 0.086 [-0.525, 0.857] |
| | Gregarious | | -0.165 [-0.296, 0.137] |
| Degree Heterogeneity | Relatively Solitary | -0.335 [-0.918, 0.319] | -0.547 [-1.128, 0.187] |
| | Gregarious | | -0.175 [-0.481, 0.078] |
| Betweeness | Relatively Solitary | -0.557 [-1.419, 0.315] | -0.403, [-1.369, 0.576] |
| | Gregarious | | 0.268 [-0.192, 0.595] |
| Degree Assortativity | Relatively Solitary | -0.429 [-1.658, 0.370] | [-0.059, 0.080] |
| | Gregarious | | **-0.552 [-0.922, -0.038]** |
| Diameter | Relatively Solitary | -0.086 [-0.662, 0.680] | -0.260 [-1.092, 0.348] |
| | Gregarious | | -0.318 [-0.590, 0.017] |
| Clustering | Relatively Solitary | 0.250 [-0.533, 1.075] | -0.096 [-0.992, 0.754] |
| | Gregarious | | **-0.466 [-0.766, -0.103]** |
| Cohesion | Relatively Solitary | -0.206 [-1.275, 0.795] | -0.825 [-1.933, 0.303] |
| | Gregarious | | **-0.443 [-0.996, -0.038]** |
| Fragmentation | Relatively Solitary | 0.385 [-0.384, 0.993] | -0.217 [-1.101, 0.404] |
| | Gregarious | | **-0.594 [-0.933, -0.313]** |

1

Figure S3: Drop lowest 5% edgeweights

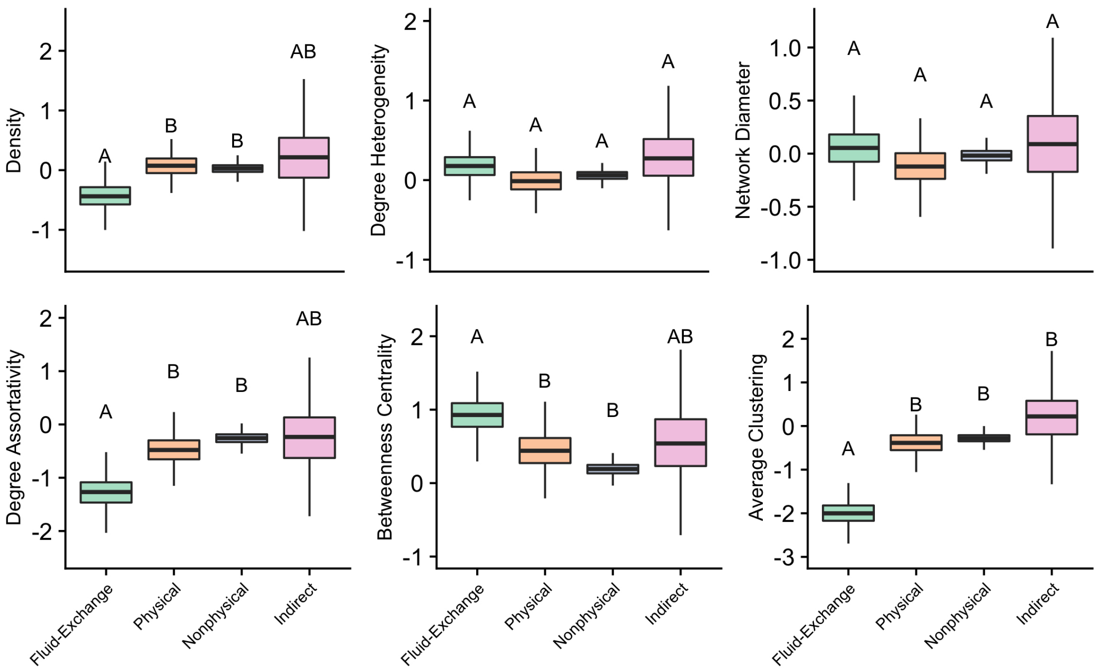


1

Figure S4: Drop lowest 15% edgeweights

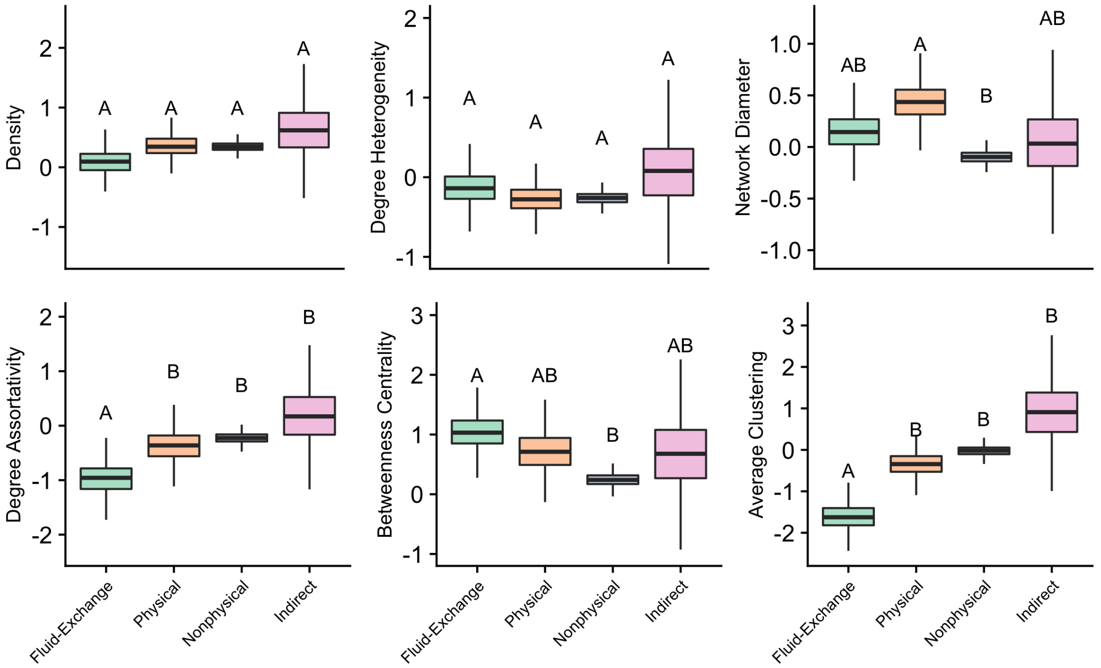


Figure S5: The predicted distributions for each network metric for contact networks in each transmission category, when the lowest 5% (a) and lowest 15% (b) of edge weights are dropped from the network. Letters represent significant differences between transmission modes.
